# Stochastic Biophysics of Cellular Radiosensitivity: From Molecular Noise and Repair Kinetics to Evolutionary Demographics

**DOI:** 10.64898/2026.08.30.748070

**Authors:** Metehan Kara, Murat Tuğrul

## Abstract

Radiation-induced DNA double-strand breaks (DSBs) drive cellular mortality, mutagenesis, and evolutionary bottlenecks. Classical phenomenological models such as the Linear-Quadratic (LQ) framework predict macroscopic survival but obscure the single-cell stochasticity that governs rare outcomes, including tumor recurrence and radioresistant persistence. We develop a stochastic differential equation framework describing DSB induction and repair as a Feller square-root process. Exact closed-form expressions for the foci mean and variance enable efficient likelihood-based inference of repair kinetics and effective molecular noise from single-cell *γ*-H2AX data without repeated Monte Carlo simulation. Coupling these kinetics to a cumulative damage hazard through the Feynman-Kac formalism links microscopic damage dynamics to macroscopic survival and provides a mechanistic mapping to the classical LQ form. Sensitivity analysis further shows that physical damage induction acts additively, whereas survival depends nonlinearly on repair rate, damage hazard, and stochastic noise. Increased noise enhances population survival by broadening the distribution of cumulative damage, consistent with Jensen’s inequality and a bet-hedging-like survival advantage for cells experiencing transiently low damage loads. This framework therefore connects stochastic single-cell biophysics with population-level radiosensitivity and provides a tractable basis for studying how cellular heterogeneity shapes survival under radiation stress.

## 1 Introduction

Ionizing radiation induces a spectrum of cellular damage, most critically DNA double-strand breaks (DSBs). Unrepaired or misrepaired DSBs can drive genomic instability, cell death, and carcinogenesis [Reya et al., 2001, Obe and Durante, 2010, Sage and Shikazono, 2017]. Understanding DSB dynamics is therefore important across evolutionary and clinical contexts, from antimicrobial resistance to cancer radiotherapy [Matsuno et al., 2021].

DSB formation and repair are inherently stochastic, generating substantial cell-to-cell heterogeneity. DSBs arise both directly from radiation and from endogenous processes including reactive oxygen species and topoisomerase activity, which modulate intrinsic radiosensitivity [Vilenchik and Knudson, 2003, Wang et al., 2019]. Repair further involves probabilistic cell-cycle checkpoints and alternative repair pathways [Houtgraaf et al., 2006, Chapman et al., 2012]. Such heterogeneity can be consequential: survival of rare cancer stem cells can contribute to tumor recurrence [Reya et al., 2001], while persistent microbial cells accumulating DSB-induced mutations can promote adaptive radioresistance [Kohanski et al., 2010, Bruckbauer et al., 2019]. Connecting these stochastic cellular dynamics to population outcomes therefore requires models that resolve single-cell variability.

Existing mathematical approaches face a trade-off between biophysical detail and computational tractability [Bodgi et al., 2016]. The Linear-Quadratic (LQ) model reliably describes macroscopic survival but does not explicitly resolve cellular stochasticity [McMahon, 2018, Wang, 2010]. Deterministic kinetic models can connect DNA repair to LQ survival or mean foci resolution [Curtis, 1986, Bodgi et al., 2016, Mariotti et al., 2013], but average over cell-to-cell variation. Conversely, detailed Monte Carlo track-structure simulations provide high biophysical resolution [Shin et al., 2021] but can be computationally demanding for parameter inference and population-level analyses [Tomezak et al., 2016, McMahon and Prise, 2019].

Here, we develop a stochastic differential equation (SDE) framework linking radiation-induced DSB dynamics to cellular survival. We model *γ*-H2AX foci as a Feller square-root process and consider both single-rate and biphasic repair. Exact analytical moments enable efficient likelihood-based calibration against single-cell *γ*-H2AX distributions without repeated Monte Carlo simulations. We calibrate the framework using experimental single-cell data and validate the inferred radiation-response kinetics against an independent dataset. We then connect these microscopic dynamics to macroscopic survival through a cumulative damage hazard and use sensitivity analysis to determine how damage induction, repair, and molecular noise shape population survival.

## 2 Theory

### 2.1 Formulation of Stochastic Foci Dynamics

The observable response to DNA double-strand breaks is not the instantaneous physical break itself, but the formation of *γ*-H2AX foci. We approximate these discrete events using a continuous stochastic differential equation (SDE), adopting a Feller square-root process [Feller, 1951, Cox et al., 1985]:

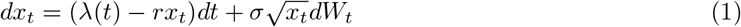

where *λ*(*t*) is the foci maturation rate, *r* the repair rate, *σ* the noise strength, and *dW*_*t*_ a standard Wiener process. The square-root noise preserves non-negativity and captures finite-count fluctuations whose variance scales with the mean.

A crucial distinction in our model is separating the instantaneous physical induction of DSBs from the delayed biological signaling response. Radiation instantaneously induces a pool of physical, unmarked DSBs, which we define using the linear-quadratic (LQ) interaction formalism [McMahon, 2018]:

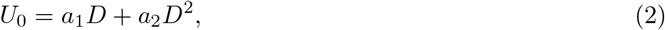

where *D* is the absorbed dose (Gy), and *a*_1_*D* and *a*_2_*D*^2^ represent single- and two-track damage contributions, respectively.

These breaks become observable as *γ*-H2AX foci following phosphorylation. Assuming first-order maturation with rate *k* gives *U* (*t*) = *U*_0_*e*^*−kt*^ and a corresponding foci flux *kU* (*t*). Including a spontaneous basal rate *a*_0_, the total foci induction rate is

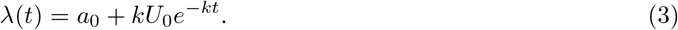

This two-state formulation captures simultaneous foci maturation and repair, including the observed peak around 30 minutes post-irradiation [Löbrich et al., 2010, Mlynarczyk et al., 2022].

We first treat the repair coefficient, *r*, as a first-order kinetic rate, wherein the clearance of DSBs scales linearly with the current damage burden [Curtis, 1986]. This assumption effectively captures the macroscopic average of the cellular capacity to resolve structural lesions via active enzymatic pathways (such as non-homologous end joining or homologous recombination). While this single-rate formulation forms the analytical core of our framework, we also relax this assumption to incorporate biphasic repair rates, accommodating the distinct fast and slow kinetics often observed in complex lesion resolution.

### 2.2 Temporal Evolution of Foci Mean and Variance

To understand the population-level dynamics prior to the onset of cell death, we can derive the exact analytical moments of the foci distribution. We evaluate the time evolution of the mean, *µ*(*t*) = E [*x*_*t*_], and the variance, *V* (*t*) = Var(*x*_*t*_), over the entire population.

Taking the expectation of the stochastic differential equation yields a deterministic ordinary differential equation (ODE) for the mean:

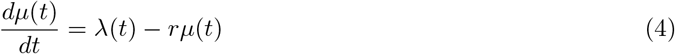

While it is mathematically convenient to assume the system rests at a perfect steady-state equilibrium prior to irradiation to reduce the parameter space, we explicitly retain the initial damage state *µ*(*t* = 0) = *x*_0_ as a free parameter. Real-world cell populations often carry transient stress loads prior to irradiation due to handling or cell-cycle positioning. Retaining *x*_0_ allows the framework to perform domain adaptation across laboratories by adjusting to specific baseline damage levels without distorting the universal radiation kinetics.

Integrating this linear ODE with our two-state maturation induction rate (Eq. 3) and free initial condition provides the exact analytical solution for the mean foci trajectory:

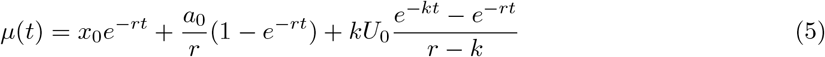

This closed-form solution mathematically separates the decay of the initial condition, the steady-state basal equilibrium, and the transient radiation-induced maturation kinetics.

A hallmark of the Feller square-root noise term 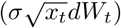 is that the variance is explicitly coupled to the mean, accurately reflecting the characteristic of Markovian birth-death processes [Feller, 1951, Cox et al., 1985]. Using Itô’s calculus, the ODE for the variance is derived as

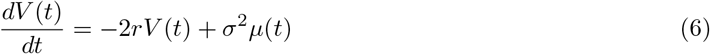

This indicates that the stochastic noise generated by the repair complexes scales dynamically with the active damage load. By solving this linear ODE using an integrating factor, the exact analytical variance can be calculated by integrating the mean trajectory:

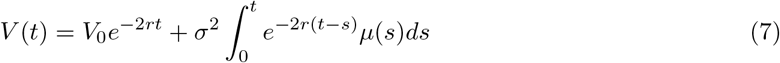

where 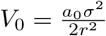 represents the baseline steady-state variance. By substituting the analytical expression for the mean (*µ*(*t*)) into the variance integral, we evaluate the time-dependent variance exactly.

Decomposing the integral into its three distinct physical components (i.e., the steady-state baseline, the transient shift from the initial experimental state *x*_0_, and the radiation-induced transient) yields the complete, closed-form analytical solution for the variance at any time *t* (see Supplementary Information for full derivation):

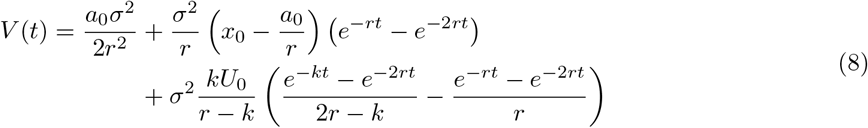

In this closed form, the fundamental dynamics become apparent: in the absence of external perturbations, the exponential terms cancel, maintaining a constant basal variance of 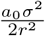. Any deviation from equilibrium or introduction of radiation damage drives a transient expansion of demographic noise that ultimately decays back to this baseline at a rate dominated by 2*r*.

To rigorously validate this analytical formulation, we checked the closed-form exact moments against large-scale Monte Carlo forward simulations of the underlying stochastic differential equation (Eq. 1). Using biologically representative parameters derived from our subsequent empirical calibration, the ensemble averages of the stochastic trajectories converge precisely to our exact expressions for both the mean and variance across all relevant timescales (**Figure S1**).

### 2.3 Maximum Likelihood Estimation (MLE) Framework

The derivation of the exact closed-form expressions for the mean (*µ*(*t*)) and variance (*V* (*t*)) represents a significant computational advantage. It permits the rapid calibration of the stochastic model using a moment-based Maximum Likelihood Estimation (MLE) against experimental single-cell data, bypassing the need for computationally exhaustive Monte Carlo forward-simulations during parameter optimization.

Let Θ = *{x*_0_, *a*_0_, *a*_1_, *a*_2_, *k, r, σ}* represent the complete set of biophysical parameters to be estimated. For a given experimental condition *j*, defined by a specific radiation dose *D*_*j*_ and post-irradiation incubation time *t*_*j*_, we observe discrete foci counts *x*_*i,j*_ across *N*_*j*_ individual cells.

Given that the Feller square-root process naturally scales variance with the mean, and assuming foci counts are sufficiently large, the state of the cell population at any time *t*_*j*_ can be approximated by a continuous Gaussian probability density function parameterized by our exact analytical moments:

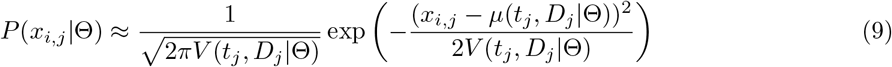

To estimate the optimal parameter set, we define the global log-likelihood function, ℒ (Θ), by summing the log-probabilities over all *i* cells across all *j* experimental conditions (doses and time points):

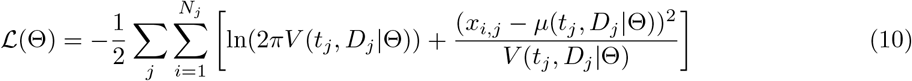

The optimal parameter set 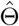 is then found by minimizing the negative log-likelihood, −ℒ (Θ) with given experimental dataset.

Crucially, formulating the likelihood using both the first and second moments simultaneously allows the framework to disentangle two coupled phenomena: the deterministic repair kinetics (*r*) and the intrinsic biological noise (*σ*). Rather than treating population variance as mere measurement error, this moment-based likelihood framework uses the observed single-cell heterogeneity to jointly infer repair kinetics and the effective stochasticity parameter.

### 2.4 Macroscopic Survival Probability

To connect our single-cell foci dynamics to macroscopic clonogenic survival, we define cell death as a cumulative Poisson process where the instantaneous hazard rate is proportional to the visible damage burden, *h*(*t*) = *c x*(*t*). The survival probability of a single cell up to time *t* is mathematically given by the expectation over all possible stochastic damage trajectories:

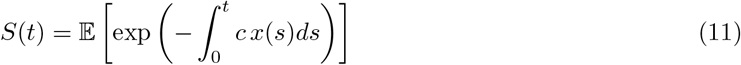

Evaluating this path integral for the Feller square-root stochastic process requires mapping the problem to a backward parabolic partial differential equation via the Feynman–Kac theorem. Because the drift and diffusion coefficients in our model are affine, the survival function admits an exact exponential-affine solution *S*(*t,x*) = exp(*−A*(*t*) *−B*(*t*)*x*) governed by a separable Riccati differential equation (see Supplementary Information for the full mathematical derivation).

For clonogenic assays, we are interested in the asymptotic surviving fraction *S*(*∞*) following an acute radiation exposure. The total radiation-induced damage pool, *U*_0_ = *a*_1_*D* + *a*_2_*D*^2^, integrates fully into the system over time, reducing the ultimate survival probability to:

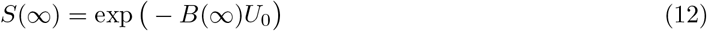

where *B*(*∞*) represents the effective biological penalty per unit of damage at steady state. By setting the time derivative of the Riccati equation to zero (*t → ∞*), we find that *B*(*∞*) must satisfy the algebraic quadratic condition:

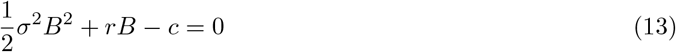

Selecting the positive physical root yields the asymptotic survival fraction:

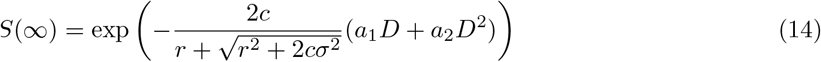

This analytical result explicitly demonstrates how stochastic noise (*σ*) fundamentally alters the macroscopic survival curve. Increasing *σ* reduces the effective damage penalty and thereby increases the surviving fraction.

Crucially, in the deterministic mean-field limit where molecular noise is absent (*σ →*0), the denominator simplifies to 2*r*, and our stochastic formulation perfectly recovers the classic macroscopic Linear–Quadratic (LQ) survival model:

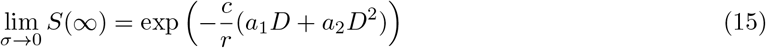

### 2.5 Extension to Biphasic Repair Kinetics

When single-compartment models fail to capture persistent or long-term tails in DNA repair data (such as unhealed damage lingering past 24 hours), it indicates that cellular repair operates via distinct pathways. To account for this, we extend our framework to a biphasic repair model. We assume that the total cellular foci pool *x*_*t*_ comprises two distinct subpopulations: a fast repair fraction (*x*_*f,t*_) governed by clearance rate *r*_*f*_, and a slow or complex repair fraction (*x*_*s,t*_) governed by clearance rate *r*_*s*_.

Newly induced DNA damage entering the system via the maturation induction rate *λ*_*t*_ = *a*_0_ + *kU*_*t*_ is partitioned into these two populations via a binomial probability *p*. Following Itô calculus for independent stochastic birth-death processes with Feller-type noise, the multi-rate SDE foundation is given by:

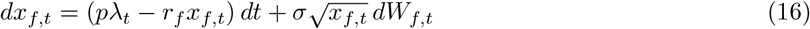

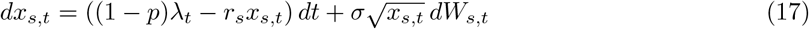

where *dW*_*f,t*_ and *dW*_*s,t*_ represent independent standard Wiener processes, and *σ* scales the intrinsic stochasticity. Total observable foci are given by the sum *x*_*t*_ = *x*_*f,t*_ + *x*_*s,t*_.

Because the fast and slow compartments evolve via independent stochastic differential equations, their respective probability distributions are mutually independent. Consequently, the mean total foci trajectory *µ*(*t*) = E [*x*_*t*_] is obtained directly by summing the individual compartment means (*µ*(*t*) = *µ*_*f*_ (*t*) + *µ*_*s*_(*t*)), and the total variance *V* (*t*) = Var(*x*_*t*_) is given by the sum of their individual variances (*V* (*t*) = *V*_*f*_ (*t*) + *V*_*s*_(*t*)). Incorporating initial conditions *x*_*f*,0_ = *px*_0_ and *x*_*s*,0_ = (1 *− p*)*x*_0_, the exact aggregate mean trajectory is given by summing the exact single-rate solutions for each compartment:

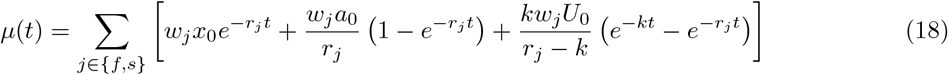

where *w*_*f*_ = *p* and *w*_*s*_ = 1 *− p*.

Similarly, rather than relying on an effective aggregate rate approximation, the exact biphasic variance is obtained by summing the independent moment equations (*V* (*t*) = *V*_*f*_ (*t*) + *V*_*s*_(*t*)), yielding the closed-form biphasic variance expression:

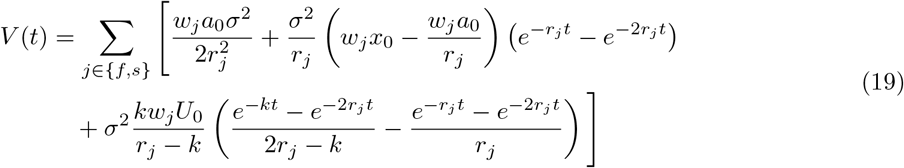

#### Maximum Likelihood Estimation (MLE) Framework for the Biphasic Model

To optimize the expanded parameter set Θ = *{x*_0_, *a*_0_, *a*_1_, *a*_2_, *k, r*_*f*_, *r*_*s*_, *p, σ}*, we utilize the exact analytical moments within a Gaussian Maximum Likelihood Estimation framework. For each experimental condition *j* defined by dose *D*_*j*_ and time *t*_*j*_, we observe discrete foci counts *x*_*i,j*_ across *N*_*j*_ cells. Assuming the Feller variance *V* (*t*_*j*_, *D*_*j*_ |Θ) properly captures single-cell heteroscedasticity, the global log-likelihood function *ℒ* (Θ) is constructed as exactly as in Eq. 10. By minimizing *ℒ* (Θ) via a global optimization, this biphasic analytical formulation simultaneously calibrates fast enzymatic clearing and slow persistent damage recovery without requiring computationally intensive Monte Carlo trajectories.

#### Analytical Derivation of Macroscopic Biphasic Survival

The stochastic survival formulation derived via the Feynman–Kac formalism extends naturally to the biphasic system. Because the fast and slow compartments are independent, the total survival path integral factorizes perfectly into the product of their independent survival expectations:

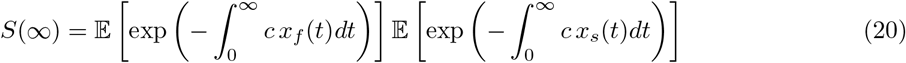

The total initial physical damage *U*_0_ = *a*_1_*D* + *a*_2_*D*^2^ is distributed to the fast and slow compartments as *pU*_0_ and (1 *− p*)*U*_0_, respectively. Evaluating the asymptotic limit for each independent compartment yields its specific steady-state penalty term *B*_*j*_(*∞*):

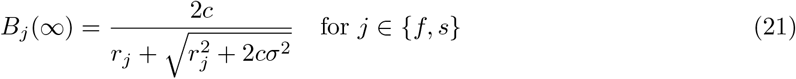

Multiplying the independent survival probabilities (adding their exponents) gives the exact closed-form clonogenic survival expression for the biphasic stochastic model:

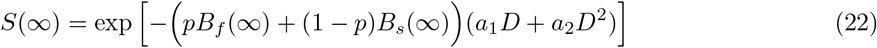

This formulation preserves the macroscopic Linear–Quadratic shape while rigorously embedding both the deterministic kinetics of partitioned double-strand break repair and the biological penalty of intrinsic molecular noise.

## 3 Results

### 3.1 Empirical Calibration and Validation of Stochastic Foci Dynamics

We used *γ*-H2AX foci as an empirical measure of the cellular DSB response, which relies on the rapid phosphorylation of histone H2AX at serine 139, which is a highly conserved molecular signaling event that initiates the cellular DSB repair cascade following exposure to ionizing radiation [Ivashkevich et al., 2012]. Because datasets reporting raw cell-by-cell foci counts across multiple doses and time points are scarce, we analyzed two studies providing single-cell *γ*-H2AX counts from irradiated human peripheral blood lymphocytes: Młynarczyk et al. [2022] for primary calibration and internal validation, and Horn et al. [2011] for external validation.

In our analysis pipeline, the Młynarczyk et al. [2022] dataset served as the primary cohort for empirical parameter calibration. Both this primary dataset and the external validation dataset explicitly include unexposed control cohorts (0 Gy), enabling estimation of the basal induction rate (*a*_0_). To reduce the parameter space, we constrained *x*_0_ to the pre-irradiation steady-state equilibrium (e.g., *x*_0_ = *a*_0_*/r* for the single-rate model). Furthermore, to maintain biological consistency, certain parameters were fixed a priori: the kinase phosphorylation rate constant was set to *k* = 6 h^*−*1^ (capturing the ~ 30-minute peak in foci formation), the *a*_1_*/a*_2_ induction ratio was constrained to 3.0 in accordance with empirical estimates [Nakamura et al., 1990], and the fast-repair fraction was established at *p* = 0.75 to account for the predominant subpopulation of rapidly repaired damage sites [Ricoul et al., 2019]. We optimized the full biophysical parameter set Θ by minimizing the negative log-likelihood of our exact analytical moments against the empirical single-cell foci data, utilizing a global Differential Evolution algorithm implemented in R [Mullen et al., 2011]. Crucially, because our Maximum Likelihood Estimation (MLE) framework natively incorporates the Feller square-root variance, we were able to extract both the deterministic repair rates (*r*) and the intrinsic molecular noise (*σ*) directly from the single-cell data (**Table 1**).

**Table 1:** Optimized Biophysical Parameters for Stochastic DSB Repair Models. Core kinetic parameters were calibrated via Maximum Likelihood Estimation against the primary single-cell dataset (Młynarczyk et al. 2022). Fixed parameters derived from biological constraints are indicated by ^*∗*^. The domain adaptation row indicates the shifted baseline parameters required to translate the model to the external dataset (Horn et al. 2011).

| Parameter | Biological Description | Single-Rate | Biphasic |
| --- | --- | --- | --- |
| **Core Kinetics (Mlynarczyk et al. Dataset)** |  |  |  |
| $a_0$ | Basal background foci induction rate (foci/h) | 0.650 | 0.449 |
| $x_0$ | Initial foci burden at $t = 0$ (foci/cell) | 1.528 | 2.860 |
| $a_1$ | Single-track physical DSB induction (breaks/Gy) | 7.918 | 6.255 |
| $a_1/a_2$ | Physical Linear-Quadratic ratio (Gy) | 3.0* | 3.0* |
| $a_2$ | Quadratic DSB induction coefficient (breaks/Gy<sup>2</sup>) | 2.639 | 2.085 |
| $k$ | Kinase maturation rate (h<sup>-1</sup>) | 6.0* | 6.0* |
| $\sigma$ | Intrinsic molecular noise scaling factor | 2.759 | 1.259 |
| $r$ | Bulk single-rate foci clearance (h<sup>-1</sup>) | 0.425 | – |
| $r_f$ | Fast compartment clearance (h<sup>-1</sup>) | – | 0.547 |
| $r_s$ | Slow compartment clearance (h<sup>-1</sup>) | – | 0.050* |
| $p$ | Proportion of damage in fast compartment | – | 0.75* |
| **Domain Adaptation (External Horn et al. Dataset)** |  |  |  |
| $a_{0,\text{Horn}}$ | Adapted background foci induction rate (foci/h) | 0.564 | 0.004 |
| $x_{0,\text{Horn}}$ | Adapted initial foci burden at $t = 0$ (foci/cell) | 1.326 | 0.025 |

For the primary dataset, which tracks foci kinetics up to 24 hours post-irradiation, we observed a profound divergence between macroscopic and microscopic evaluation metrics. If evaluated solely on macroscopic averages, the biphasic model appeared to offer a marginally tighter fit to the mean trajectory (Trend *R*^2^ = 0.9128) compared to the single-rate model (Trend *R*^2^ = 0.8330). However, when evaluating the models against the true single-cell distribution via the Akaike Information Criterion (AIC), the single-rate model demonstrated overwhelmingly superior performance to the biphasic model (ΔAIC = 5638.89; **Table 2**). This indicates that for the first 24 hours, the fundamental Feller square-root variance of the single-rate formulation captures the intrinsic biological noise of the population, whereas the added structural complexity of the biphasic model overfits the means at the expense of accurately predicting cellular heterogeneity (**Figure 1A, B**).

**Table 2:** Comprehensive Model Evaluation Metrics. Goodness-of-fit and information-theoretic metrics for the Single-Rate and Biphasic repair models across all evaluated datasets. ΔAIC is calculated independently within each dataset cohort. Trend *R*^2^ evaluates the macroscopic mean trajectory, while Single-Cell *R*^2^ reflects microscopic variance.

| Dataset Cohort | Model | $K$ | N Cells | Trend $R^2$ | Single-Cell $R^2$ | Log-Likelihood | $\Delta\text{AIC}$ |
| --- | --- | --- | --- | --- | --- | --- | --- |
| Primary Calibration<br>(Młynarczyk et al.) | Single-Rate | 4 | 29,000 | 0.8330 | 0.3291 | -93,774.94 | **0.00** |
|  | Biphasic | 5 | 29,000 | 0.9128 | 0.3713 | -96,593.39 | 5638.89 |
| Internal Validation<br>(Młynarczyk et al.) | Single-Rate | 4 | 34,000 | 0.6521 | 0.2700 | -116,679.09 | **0.00** |
|  | Biphasic | 5 | 34,000 | 0.9002 | 0.3620 | -117,532.39 | 1708.61 |
| External Validation<br>(Horn et al.) | Single-Rate | 4 | 2,000 | 0.7699 | 0.6816 | -5,785.40 | 1229.05 |
|  | Biphasic | 5 | 2,000 | 0.9034 | 0.7998 | -5,169.87 | **0.00** |

**Figure 1:**
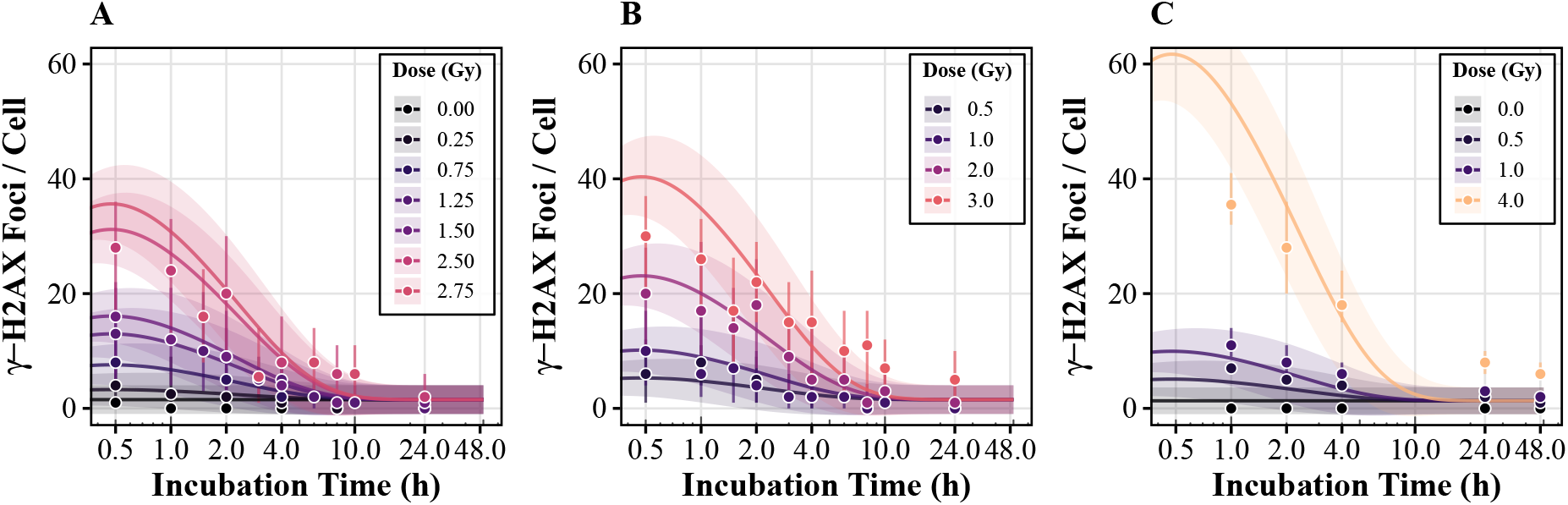
Empirical calibration and validation of the stochastic single-rate DSB repair model. Experimental single-cell *γ*-H2AX foci distributions are plotted alongside the exact analytical moments of the single-rate stochastic model. Solid lines represent the deterministic mean trajectory *µ*(*t*), while the shaded regions indicate the Feller square-root standard deviation 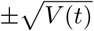. **(A)** Model calibration against the primary dataset (Młynarczyk et al., 2022), demonstrating accurate capture of initial induction, the damage peak, and 24-hour bulk clearance. **(B)** Internal validation against an independent cohort from the primary dataset, confirming robust prediction of both macroscopic kinetics and microscopic cellular heterogeneity. **(C)** External validation and domain adaptation using the Horn et al. (2011) dataset. While the single-rate framework accurately translates across laboratory environments for the acute 24-hour response, it predictably underestimates the long tail of persistent, complex damage that remains at 48 hours.

To rigorously test the universality of these stochastic kinetics, we performed an external validation against the independent Horn et al. [2011] dataset, which extends the experimental observation window to 48 hours. Real-world cell populations exhibit baseline variations due to handling stress and distinct laboratory environments, as evidenced by the distinct 0 Gy control baselines in the two datasets. To account for this without distorting the universal radiation kinetics, we performed a targeted domain adaptation: we recalibrated only the spontaneous background damage (*a*_0_) for the Horn cohort, while keeping the core radiation-induced kinetic parameters strictly anchored to the values derived from the primary Młynarczyk dataset. While the single-rate model translates remarkably well across laboratories for the first 24 hours, the extended 48-hour timeline exposes a long tail of persistent, unhealed damage that the single-rate formulation predictably underestimates (**Figure 1C**), where the biphasic model proves its necessity. By partitioning the damage into fast and slow clearing compartments, the biphasic formulation successfully resolved the 48-hour persistent damage tail, yielding decisively superior statistical support in favor of the biphasic model (ΔAIC = 1229.05; **Table 2**) (**Figure S1**).

### 3.2 From Microscopic Foci to Macroscopic Survival

As demonstrated by our empirical calibration, the theoretical framework seamlessly accommodates both bulk clearance (single-rate) and complex, long-term persistent damage (biphasic) kinetics. The biphasic extension provides a critical mathematical tool for researchers evaluating delayed damage tails beyond 24 hours. However, the majority of standard clonogenic survival assays and theoretical endpoints evaluate cellular fate based on the bulk repair achieved within the first 24 hours. This is a time regime where our single-rate model is both mathematically and statistically optimal, providing a more intuitive mapping between microscopic foci kinetics and macroscopic cellular fate. Therefore, we utilize the single-rate formulation to analyze the survival principles and conduct the sensitivity analyses. As established in the Theory section, these exact survival expressions translate identically to the biphasic formulation.

To translate these validated microscopic repair kinetics into macroscopic population demographics, we define cellular survival as a cumulative hazard process. To ensure our macroscopic survival projections are biologically relevant, we utilize the empirically estimated biophysical parameters (**Table 1**) in all subsequent theoretical analyses. In this framework, the instantaneous probability of cell death is directly proportional to the active foci number with a fundamental lethality constant *c*, i.e., *h* = *c x*_*t*_.

By integrating this hazard over the full distribution of stochastic damage trajectories, our exact analytical solution for the asymptotic surviving fraction, *S*(*∞*) = exp(*−B*(*∞*)(*a*_1_*D* + *a*_2_*D*^2^) as derived in Eq. (14), bridges the gap between single-cell biophysics and macroscopic clinical demographics. The biological penalty term is given by 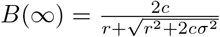 (see Theory and SI for detailed derivations). In the deterministic limit where intrinsic demographic noise is assumed to be zero (*σ →* 0), our framework perfectly recovers the classic phenomenological Linear-Quadratic (LQ) survival model, *S* = exp(*−αD −βD*^2^), yielding the direct equivalencies: 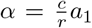 and 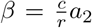. This deterministic reduction provides a powerful mechanistic interpretation of the traditional LQ parameters: macroscopic radiosensitivity is fundamentally defined by the ratio of the microscopic lethality penalty (*c*) to the repair rate (*r*), scaled by the physical induction of single- and two-track damage (*a*_1_, *a*_2_).

For the remainder of this analysis, we parameterize the specific lethality constant as *c* = 0.014 h^*−*1^foci^*−*1^ by combining our empirically calibrated single-rate kinetics (*r, a*_1_) and *α* estimates obtained from other in vitro radiosensitivity studies of human peripheral blood lymphocytes [Nakamura et al., 1990, McMahon, 2018], providing a biologically grounded anchor for our subsequent demographic and evolutionary simulations. Also, we keep spontaneous background damage to zero (*a*_0_ = 0) to isolate purely radiation-induced kinetics and mortality.

To visualize these dynamics, we performed stochastic forward-simulations of the foci trajectories following Eq. (1) and incorporating this lethality constant across multiple radiation doses (**Figure 2A**). Tracking the corresponding cumulative survival over time reveals that the survival probability descends rapidly during the initial damage maturation phase, but stabilizes as enzymatic repair successfully clears the lesion burden (**Figure 2B**). Because the characteristic repair rate (*r* = 0.425 h^*−*1^) effectively depletes the transient radiation-induced foci within a day, the survival curves reach a flat equilibrium well within the 24-hour observation window. When we plot this 24-hour surviving fraction against a broader dose range (0 to 10 Gy), the classic downward-bending Linear-Quadratic topology emerges naturally from the underlying single-cell kinetics (**Figure 2C**). Overlaid on this plot, our exact analytical prediction for the asymptotic limit *S*(*∞*) in Eq. (14) aligns perfectly with the 24-hour stochastic simulations, confirming that *t* = 24 h serves as a robust functional asymptote for clonogenic fate. Furthermore, to explicitly connect these acute radiation responses to population demographics, we evaluated the cellular lifespan statistics. As radiation dose increases, the escalating foci burden drives a severe collapse in both the mean lifespan and its intrinsic variance (**Figure 2C**, right axis), demonstrating how stochastic damage accumulation fundamentally constrains a population’s evolutionary potential.

**Figure 2:**
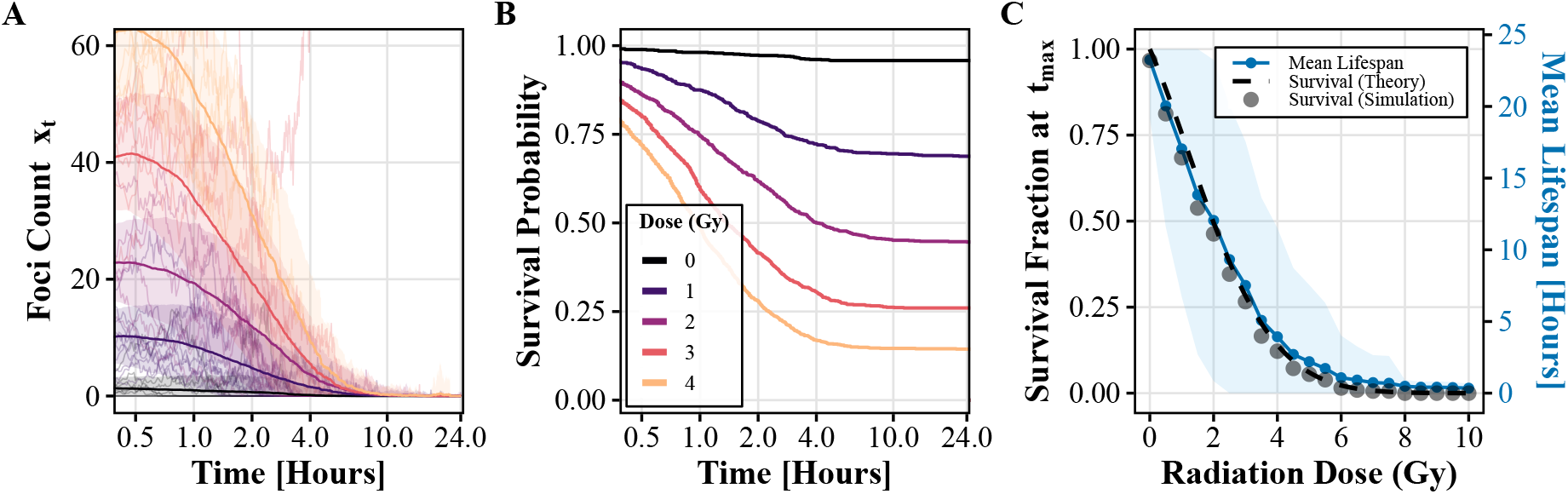
Bridging microscopic stochastic damage to macroscopic Linear-Quadratic survival and population demographics. **(A)** Stochastic single-rate model forward-simulations of single-cell *γ*-H2AX foci trajectories across acute radiation doses (*D* = 0, 1, 2, 3, 4 Gy). Simulations are driven by the empirically calibrated single-rate parameters (*r* = 0.425 h^*−*1^, *σ* = 2.759) and a cell-lethality constant *c* = 0.014 h^*−*1^foci^*−*1^. To focus on radiation based survival, we kept *a*_0_ = 0 for this analysis. **(B)** Corresponding time-dependent survival probabilities derived from the cumulative damage hazard. Survival plummets rapidly during the initial accumulation of DNA damage but reaches a stable, flat equilibrium within 24 hours as enzymatic repair clears the transient lesion burden. **(C)** The emergent macroscopic dose-response and demographic collapse (0 to 10 Gy). *Left axis:* The simulated 24-hour surviving fraction (points) perfectly matches the exact analytical prediction for the asymptotic limit *S*(*∞*) (solid line), naturally recovering the classic downward-bending Linear-Quadratic (LQ) topology entirely from underlying microscopic kinetics. *Right axis:* The mean cellular lifespan and its intrinsic variance (shaded area). As the initial foci burden escalates with dose, both the expected lifespan and its variance experience a severe demographic collapse.

### 3.3 Sensitivity Analysis of Cellular Fate

To uncover the biophysical drivers of cellular radiosensitivity and their evolutionary implications, we conducted a systematic sensitivity analysis, mapping macroscopic survival fraction against targeted perturbations of our empirically estimated kinetic parameters. By exploring how variations in these core rates dictate demographic fate, we aim to bridge single-cell repair mechanics with population-level evolutionary constraints.

We first focused on the intrinsic biological repair machinery, scaling the enzymatic repair rate (*r*) and the molecular noise (*σ*) from 0 to 200% of their calibrated baselines at a fixed acute radiation dose of *D* = 2 Gy to generate an analytical survival landscape (**Figure 3A**). To confirm the robustness of these theoretical topographies, we simultaneously performed full stochastic forward-simulations across the identical parameter space (**Figure 3B**). These simulations demonstrated good congruence with our exact analytical solutions for the asymptotic survival fraction *S*(*∞*) and the biological penalty term 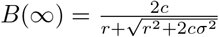 derived in the Theory section.

**Figure 3:**
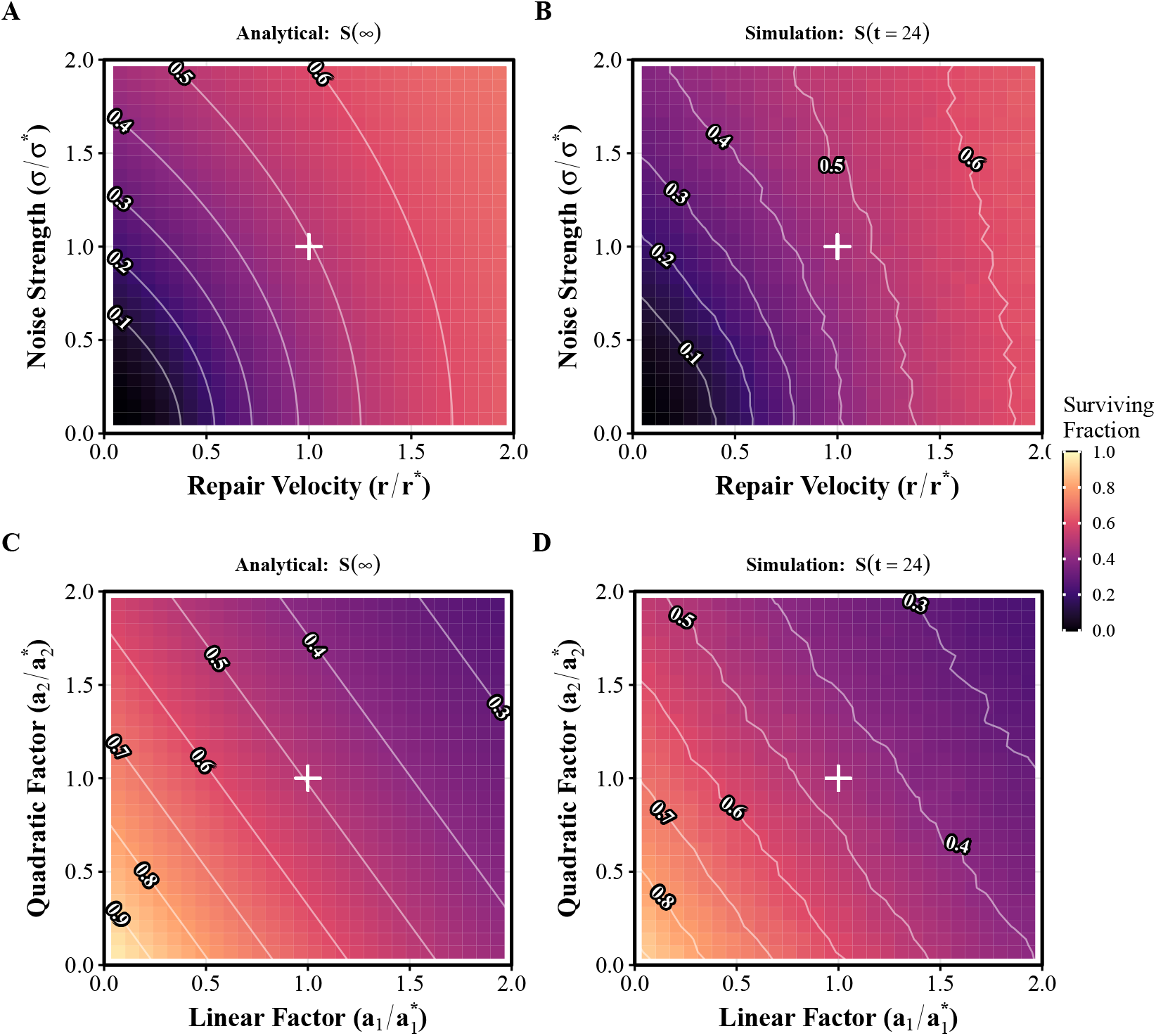
Sensitivity analysis of the biophysical parameters governing macroscopic cell survival. **(A)** Exact analytical survival fraction, *S*(*∞*), evaluated at a fixed acute dose of *D* = 2 Gy, scaling the enzymatic repair rate (*r*) and molecular noise (*σ*) from 0 to 2*×* their empirically estimated baselines. The nonlinear topography reveals diminishing survival returns with increasing repair efficiency and highlights the survival advantage associated with increased molecular noise. **(B)** Full stochastic forward-simulations corresponding to the parameter space in (A), showing good congruence with the analytical surface. **(C)** Analytical sensitivity profiling of the physical damage induction parameters: single-track (*a*_1_) and two-track (*a*_2_) coefficients, evaluated at a fixed dose of *D* = 2 Gy. The perfectly straight contour lines visually confirm the independent, additive nature of physical damage induction. **(D)** Full stochastic forward-simulations corresponding to the induction parameter space in (C), validating the robust geometric linearity.

Analyzing the enzymatic repair rates (*r*) reveals a highly asymmetric survival landscape. The empirically estimated baseline lies in a region of pronounced asymmetry in the survival response. Decreasing the repair capacity by 50% (0.5*×*) causes a severe collapse in clonogenic viability, plunging the survival fraction from approximately 0.5 down to around 0.3. In contrast, increasing the repair capacity by 50% (1.5 *×*) yields strictly diminishing returns, elevating survival to only roughly 0.6. Investing excess cellular energy to further accelerate repair offers marginal fitness gains compared to the catastrophic demographic penalty of falling below the baseline threshold.

The molecular noise parameter (*σ*) presents an even more intriguing evolutionary dynamic. At the baseline repair rate, suppressing the noise entirely toward the deterministic limit (*σ →* 0) exerts negligible impact on the overall survival. Conversely, amplifying the intrinsic noise strictly increases the surviving fraction by 10 *−* 20%. Furthermore, the fitness topography reveals a critical interaction between these two axes: if the repair rate is significantly impaired (low *r*), the population’s survival becomes highly fragile to decreases in noise. In this repair-deficient regime, the gradient along the *σ* axis becomes remarkably steep, indicating that the survival benefit associated with stochastic heterogeneity becomes particularly pronounced. Because the lethality hazard scales exponentially with the damage burden, widening the phenotypic distribution via elevated molecular noise (*σ*) increases the probability of cells experiencing lower-than-average damage trajectories. Thus, rather than functioning as an inefficient biological error, intrinsic molecular noise acts as a robust population-level buffer against acute evolutionary bottlenecks.

Finally, we varied the physical damage induction parameters (*a*_1_, *a*_2_; **Figure 3C**). Similarly, our exact analytical predictions for this induction landscape show good agreement with the full stochastic simulations (**Figure 3D**). In contrast to the nonlinear interaction between *r* and *σ*, the induction landscape exhibits straight contours, reflecting the additive dependence *a*_1_*D* + *a*_2_*D*^2^ in the survival expression. At *D* = 2 Gy, survival is more sensitive to changes in *a*_1_ than *a*_2_. Thus, physical damage induction enters additively, whereas repair and stochasticity jointly determine the nonlinear biological survival response.

## 4 Discussion

We developed a stochastic differential equation (SDE) framework linking radiation-induced DNA damage at the single-cell level to population survival. By deriving closed-form expressions for the mean (*µ*(*t*)) and variance (*V* (*t*)) of *γ*-H2AX foci dynamics, the framework enables efficient likelihood-based inference of repair kinetics and cellular stochasticity without repeated Monte Carlo simulations. Importantly, these dynamics were calibrated against single-cell experimental data and validated against an independent dataset, providing an empirically grounded bridge between microscopic damage-repair processes and macroscopic survival.

A key consequence of this framework is a mechanistic mapping to the classical Linear–Quadratic (LQ) survival model. Given the physical damage induction *U*_0_ = *a*_1_*D* + *a*_2_*D*^2^, integration of the stochastic damage dynamics through a cumulative death hazard preserves the LQ survival form. In the deterministic limit (*σ →* 0), the model yields *S* = exp(*− αD−βD*^2^), with the effective radiosensitivity parameters determined by microscopic damage induction, repair and lethality. Parameterization using established in vitro radiosensitivity estimates for human lymphocytes further anchors this mapping to experimentally relevant regimes [Nakamura et al., 1990, McMahon, 2018].

Beyond this micro-to-macro mapping, our sensitivity analysis reveals a marked distinction between physical damage induction and the subsequent biological response. The induction parameters (*a*_1_, *a*_2_) contribute additively to the initial damage burden, whereas survival depends nonlinearly on repair kinetics (*r*) and molecular noise (*σ*). The calibrated repair rate lies in a region where reductions in repair produce substantially greater survival losses than the gains obtained from comparable increases in repair. Such diminishing returns could be consistent with evolutionary trade-offs if faster repair carries metabolic or physiological costs.

A central result is that molecular noise can itself enhance population survival. The Feynman–Kac survival formulation shows that increasing *σ* reduces the effective damage penalty, while the convexity of the exponential survival function provides an intuitive explanation through Jensen’s inequality: stochastic heterogeneity generates cells experiencing lower-than-average cumulative damage and therefore disproportionately contributing to the surviving population. This produces a bet-hedging-like survival advantage without requiring genetic heterogeneity. Such stochastic variation may therefore buffer populations during severe evolutionary bottlenecks, consistent with broader roles proposed for phenotypic variability in evolutionary dynamics [Weinreich et al., 2026].

Several limitations suggest natural extensions. The available *γ*-H2AX datasets provide single-cell distributions at fixed time points rather than longitudinal trajectories of individual cells [Horn et al., 2011, Młynarczyk et al., 2022]; real-time imaging could therefore provide stronger tests of the predicted stochastic dynamics [Robert et al., 2018]. Moreover, the Feller diffusion approximates discrete molecular events continuously, whereas low-copy-number dynamics or cell divisions may require jump-process formulations [Tuğrul and Steiner, 2025]. Finally, incorporating track-structure microdosimetry would allow the framework to distinguish radiation qualities and clustered versus sparse DNA damage [Nakano et al., 2022].

Overall, this framework provides a tractable bridge between stochastic cellular damage-repair dynamics and population survival, with applications ranging from radiobiology to the evolutionary consequences of non-genetic cellular heterogeneity.

## Acknowledgements

M.T. was supported by the European Union’s Marie Skłodowska-Curie Actions (Grant 101069035) and Horizon 2020 European Research Council (Grant 864971); M.K. by the KYK Scholarship of the Ministry of Youth and Sports of the Republic of Türkiye. We thank Dr. Simone Mörtl (TUM) for discussions on the biological background and Dr. Stephen Barnard (UKHSA) for providing the published *γ*-H2AX dataset of Horn et al. (2011).

## Supplementary Information

### Derivation of the Variance Ordinary Differential Equation

Let the stochastic foci dynamics be governed by the following Stochastic Differential Equation (SDE):

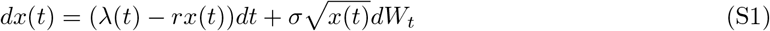

where *µ*(*t*) = E[*x*(*t*)] and *V* (*t*) = Var(*x*(*t*)). Taking the expectation of the SDE yields the ordinary differential equation for the mean:

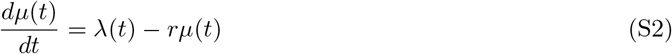

To find the deterministic evolution of the variance, we first determine the expectation of the second moment, E [*x*(*t*)^2^]. Applying Itô’s Lemma to the function *f* (*x*) = *x*^2^, we obtain:

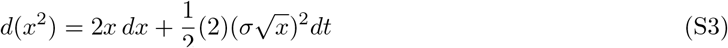

Substituting the original SDE for *dx*:

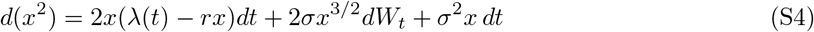

Taking the expectation on both sides (noting that the expectation of the Itô integral with respect to *dW*_*t*_ is zero) yields the ODE for the second moment:

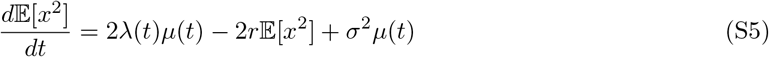

By definition, the variance is *V* (*t*) = E [*x*^2^] *−µ*(*t*)^2^. Taking the time derivative of this definition and applying the chain rule gives:

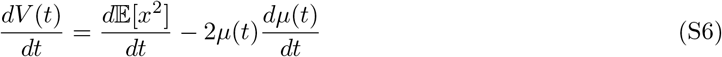

Substituting Equation S2 and Equation S5 into this expression:

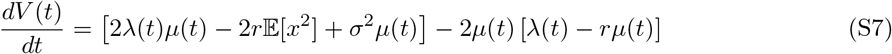

Expanding the terms:

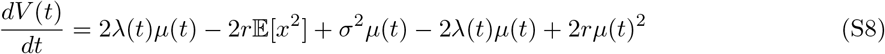

The deterministic source terms 2*λ*(*t*)*µ*(*t*) perfectly cancel. Grouping the remaining terms by *−*2*r* yields:

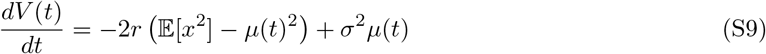

Which simplifies exactly to the final variance ODE:

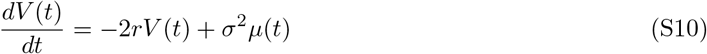

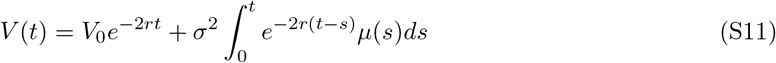

where 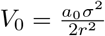 represents the baseline steady-state variance.

By substituting the analytical expression for the mean (*µ*) into the variance integral, we can evaluate the time-dependent variance exactly:

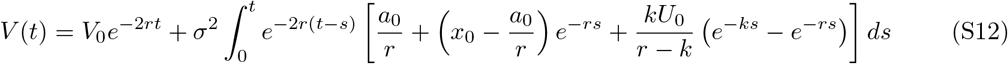

The integral can be decomposed into three distinct components. Evaluating the baseline steady-state component gives:

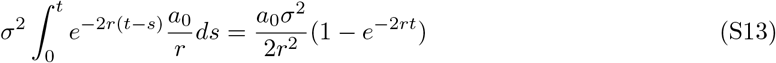

Since the initial baseline variance is 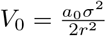, the *e*^*−*2*rt*^ terms cancel out, demonstrating that the basal variance remains constant at 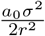 over time in the absence of external perturbations.

Evaluating the transient shift caused when the experimental initial state *x*_0_ deviates from perfect equilibrium yields:

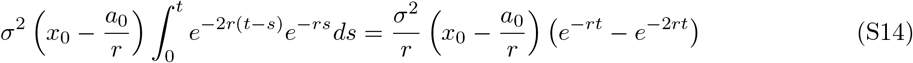

Finally, evaluating the radiation-induced transient component yields:

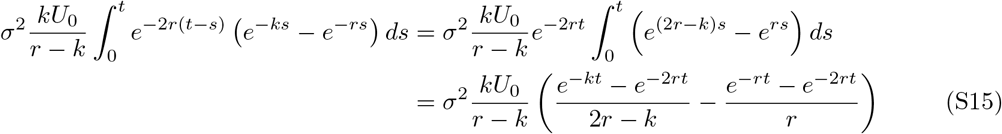

Combining these terms provides the complete, closed-form analytical solution for the variance of the visible foci at any time *t*:

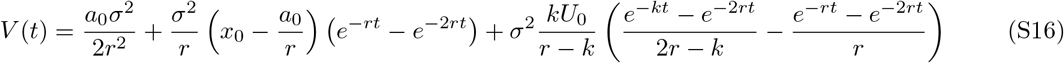

### Derivation of Exact Asymptotic Survival via the Feynman–Kac Formalism

We consider the stochastic dynamics of visible damage foci *x*(*t*) governed by the Feller square-root process:

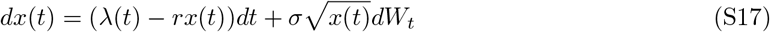

The survival probability of a single cell up to time *t*, given an initial foci state *x*(0) = *x*_0_, under the hazard function *h*(*t*) = *c x*(*t*) is defined as the expectation of the path integral:

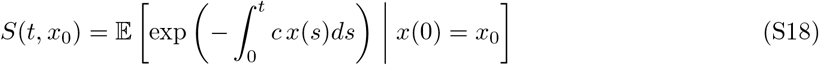

#### The Feynman–Kac Partial Differential Equation

By the Feynman–Kac theorem, the function *S*(*t, x*) satisfies the backward parabolic partial differential equation:

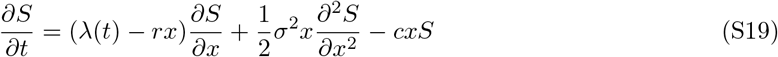

subject to the initial condition *S*(0, *x*) = 1.

Because the drift and diffusion coefficients of the square-root process are affine in *x*, we adopt the standard exponential-affine ansatz from the Cox–Ingersoll–Ross (CIR) framework:

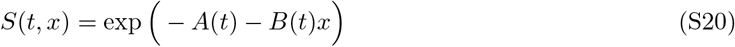

with the initial boundary conditions *A*(0) = 0 and *B*(0) = 0.

Computing the partial derivatives of the ansatz:

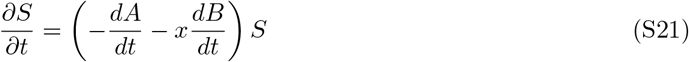

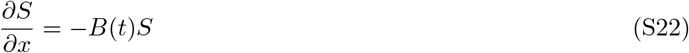

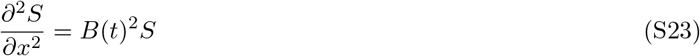

Substituting these into Equation (S19) and dividing both sides by *S*(*t, x*):

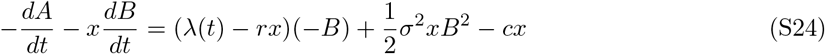

Grouping terms by powers of *x*:

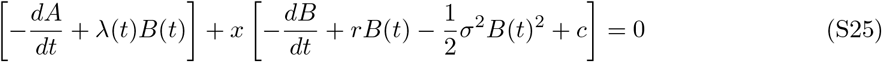

For this identity to hold for all values of *x*, both bracketed terms must vanish independently, yielding two coupled ordinary differential equations:

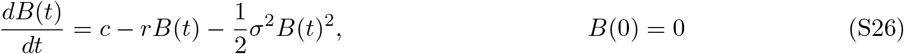

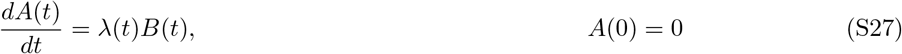

### Analytical Solution of the Riccati Equation

Equation (S26) is a separable Riccati differential equation. Rearranging:

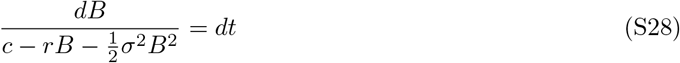

Let 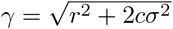. The quadratic denominator can be factored as 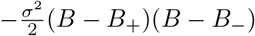, where the roots are:

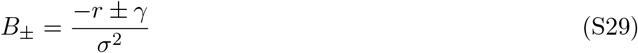

Integrating by partial fractions with the initial condition *B*(0) = 0 yields the exact time-dependent solution:

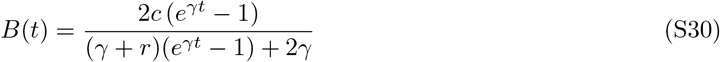

#### Asymptotic Limit and Clonogenic Survival

To determine the final surviving fraction *S*(*∞*) following an acute radiation exposure, we evaluate the system as *t → ∞*.

In the acute regime, the baseline spontaneous induction is negligible (*a*_0_ *≈* 0). The total radiation-induced damage pool *U*_0_ = *a*_1_*D* + *a*_2_*D*^2^ enters the system either instantaneously at *t* = 0 (*x*_0_ = *U*_0_) or through rapid kinase maturation (*λ*(*t*) = *kU*_0_*e*^*−kt*^). In both formulations, the cumulative total damage burden integrates to *U*_0_, reducing the asymptotic survival to:

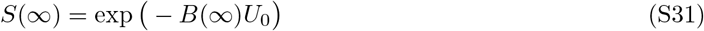

Taking the limit of Equation (S30) as *t → ∞*, the growing exponential *e*^*γt*^ dominates all constant terms:

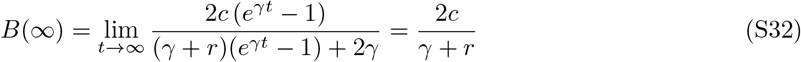

Substituting 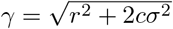 back into the expression yields:

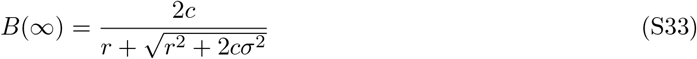

#### Alternative Stationary Proof

Alternatively, at steady state (*t → ∞*), the time derivative vanishes: 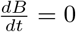. Equation (S26) reduces to the algebraic quadratic equation:

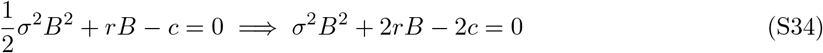

Applying the quadratic formula and selecting the positive physical root:

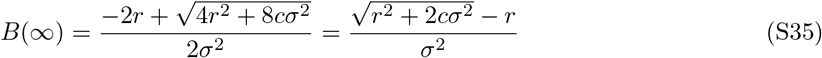

Rationalizing the numerator by multiplying numerator and denominator by 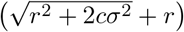:

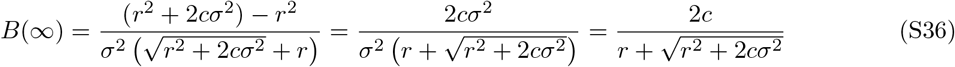

Substituting this asymptotic scaling back into the survival function with *U*_0_ = *a*_1_*D* + *a*_2_*D*^2^ yields the exact closed-form clonogenic survival expression:

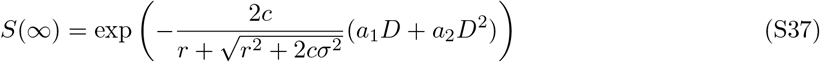

In the deterministic zero-noise limit (*σ →*0), the denominator simplifies to 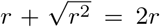, exactly recovering the classic mean-field Linear–Quadratic formulation:

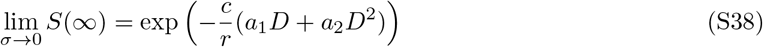

**Figure S1:**
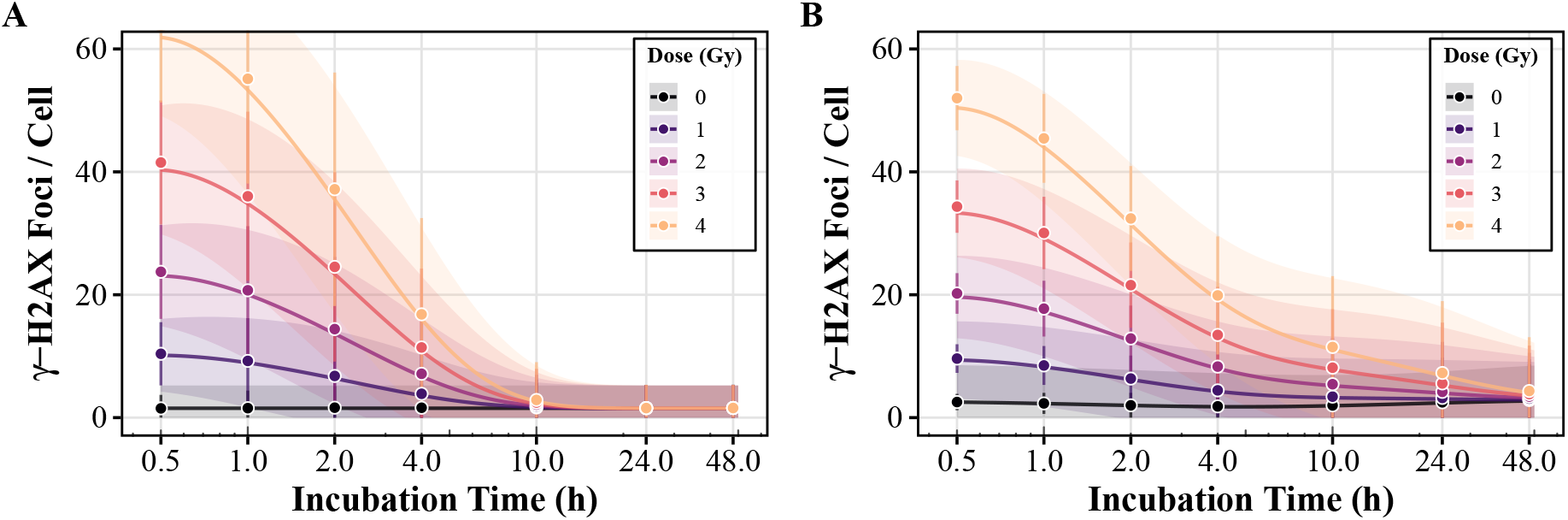
Validation of exact analytical moments against stochastic forward-simulations. Using the empirically fitted parameters from the single-rate model (**A**, corresponding to Figure 1) and the biphasic model (**B**, corresponding to Figure S2), we demonstrate that the exact analytical moments match the stochastic simulation trajectories with high precision across all relevant timescales. Solid lines represent the analytical total mean trajectory *µ*(*t*), and shaded regions indicate the analytical total standard deviation 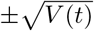. Discrete markers with error bars represent the ensemble mean and standard deviation derived from full stochastic forward-simulations. This close congruence highlights the computational superiority and robustness of the analytical results, circumventing the need for exhaustive Monte Carlo approximations.

**Figure S2:**
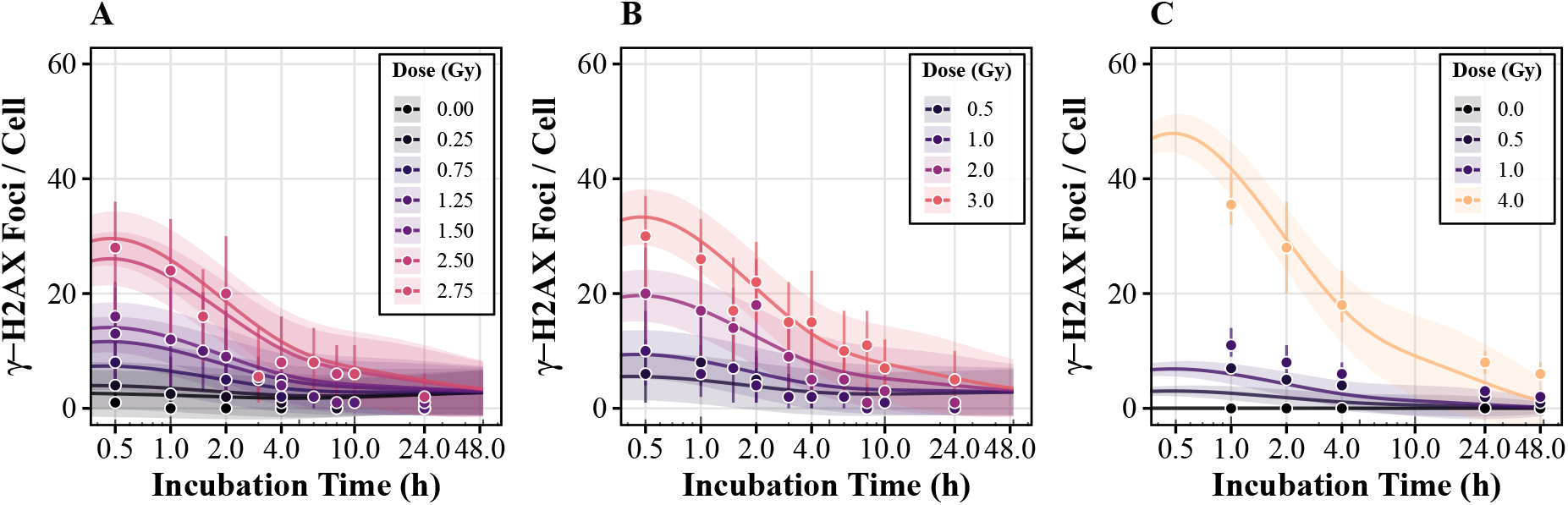
Empirical calibration and validation of the stochastic biphasic DSB repair model. Experimental single-cell *γ*-H2AX foci distributions are plotted alongside the exact analytical moments of the extended biphasic stochastic model. Solid lines represent the deterministic aggregate mean trajectory *µ*(*t*), while the shaded regions indicate the composite Feller square-root standard deviation 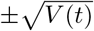. **(A)** Model calibration against the primary dataset (Młynarczyk et al., 2022). The biphasic formulation captures the initial induction and damage peak by partitioning the lesion burden into distinct fast and slow repair compartments. **(B)** Internal validation against an independent cohort from the primary dataset, mapping the 24-hour macroscopic and microscopic kinetics. **(C)** External validation and domain adaptation using the extended 48-hour dataset (Horn et al., 2011). In contrast to the single-rate framework, the biphasic model successfully resolves the long-term biological response, accurately capturing the extended tail of persistent, complex damage that remains unhealed at 48 hours.

## References

Larry Bodgi, Aurélien Canet, Laurent Pujo-Menjouet, Annick Lesne, Jean-Marc Victor, and Nicolas Foray. Mathematical models of radiation action on living cells: From the target theory to the modern approaches. A historical and critical review. Journal of Theoretical Biology, 394:93–101, April 2016. ISSN 00225193. doi: 10.1016/j.jtbi.2016.01.018.

Steven T. Bruckbauer, Joseph D. Trimarco, Joel Martin, Brian Bushnell, Katherine A. Senn, Wendy Schackwitz, Anna Lipzen, Matthew Blow, Elizabeth A. Wood, Wesley S. Culberson, Christa Pennacchio, and Michael M. Cox. Experimental Evolution of Extreme Resistance to Ionizing Radiation in Escherichia coli after 50 Cycles of Selection. Journal of Bacteriology, 201(8):10.1128/jb.00784-18, March 2019. doi: 10.1128/jb.00784-18.

J. Ross Chapman, Martin R. G. Taylor, and Simon J. Boulton. Playing the End Game: DNA DoubleStrand Break Repair Pathway Choice. Molecular Cell, 47(4):497–510, August 2012. ISSN 1097-2765. doi: 10.1016/j.molcel.2012.07.029.

John C. Cox, Jonathan E. Ingersoll, and Stephen A. Ross. A Theory of the Term Structure of Interest Rates. Econometrica, 53(2):385–407, 1985. ISSN 0012-9682. doi: 10.2307/1911242.

S. B. Curtis. Lethal and potentially lethal lesions induced by radiation–a unified repair model. Radiation Research, 106(2):252–270, May 1986. ISSN 0033-7587.

William Feller. Two Singular Diffusion Problems. Annals of Mathematics, 54(1):173–182, 1951. ISSN 0003-486X. doi: 10.2307/1969318.

Simon Horn, Stephen Barnard, and Kai Rothkamm. Gamma-H2AX-Based Dose Estimation for Whole and Partial Body Radiation Exposure. PLOS ONE, 6(9):e25113, September 2011. ISSN 1932-6203. doi: 10.1371/journal.pone.0025113.

Jaco H. Houtgraaf, Jorie Versmissen, and Wim J. van der Giessen. A concise review of DNA damage checkpoints and repair in mammalian cells. Cardiovascular Revascularization Medicine, 7(3):165–172, July 2006. ISSN 1553-8389. doi: 10.1016/j.carrev.2006.02.002.

Alesia Ivashkevich, Christophe E. Redon, Asako J. Nakamura, Roger F. Martin, and Olga A. Martin. Use of the γ-H2AX assay to monitor DNA damage and repair in translational cancer research. Cancer Letters, 327(1):123–133, December 2012. ISSN 0304-3835. doi: 10.1016/j.canlet.2011.12.025.

Michael A. Kohanski, Mark A. DePristo, and James J. Collins. Sublethal Antibiotic Treatment Leads to Multidrug Resistance via Radical-Induced Mutagenesis. Molecular Cell, 37(3):311–320, February 2010. ISSN 1097-2765. doi: 10.1016/j.molcel.2010.01.003.

Markus Löbrich, Atsushi Shibata, Andrea Beucher, Anna Fisher, Michael Ensminger, Aaron A. Goodarzi, Olivia Barton, and Penny A. Jeggo. γH2AX foci analysis for monitoring DNA double-strand break repair: Strengths, limitations and optimization. Cell Cycle, 9(4):662–669, February 2010. ISSN 1538-4101. doi: 10.4161/cc.9.4.10764.

Luca G. Mariotti, Giacomo Pirovano, Kienan I. Savage, Mihaela Ghita, Andrea Ottolenghi, Kevin M. Prise, and Giuseppe Schettino. Use of the γ-H2AX Assay to Investigate DNA Repair Dynamics Following Multiple Radiation Exposures. PLOS ONE, 8(11):e79541, November 2013. ISSN 1932-6203. doi: 10.1371/journal.pone.0079541.

Yusuke Matsuno, Mai Hyodo, Mafuka Suzuki, Yosuke Tanaka, Yasunori Horikoshi, Yasufumi Murakami, Hidetaka Torigoe, Hiroyuki Mano, Satoshi Tashiro, and Ken-ichi Yoshioka. Replication-stress-associated DSBs induced by ionizing radiation risk genomic destabilization and associated clonal evolution. iScience, 24(4), April 2021. ISSN 2589-0042. doi: 10.1016/j.isci.2021.102313.

Stephen J. McMahon and Kevin M. Prise. Mechanistic Modelling of Radiation Responses. Cancers, 11 (2):205, February 2019. ISSN 2072-6694. doi: 10.3390/cancers11020205.

Stephen Joseph McMahon. The linear quadratic model: Usage, interpretation and challenges. Physics in Medicine & Biology, 64(1):01TR01, December 2018. ISSN 0031-9155. doi: 10.1088/1361-6560/aaf26a.

Dorota Młynarczyk, Pedro Puig, Carmen Armero, Virgilio Gómez-Rubio, Joan F. Barquinero, and Mónica Pujol-Canadell. Radiation dose estimation with time-since-exposure uncertainty using the $$\gamma $$-H2AX biomarker. Scientific Reports, 12(1):19877, November 2022. ISSN 2045-2322. doi: 10.1038/s41598-022-24331-1.

Katharine M. Mullen, David Ardia, David L. Gil, Donald Windover, and James Cline. DEoptim: An R Package for Global Optimization by Differential Evolution. Journal of Statistical Software, 40:1–26, April 2011. ISSN 1548-7660. doi: 10.18637/jss.v040.i06.

N. Nakamura, Y. Kusunoki, and M. Akiyama. Radiosensitivity of CD4 or CD8 positive human T-lymphocytes by an in vitro colony formation assay. Radiation Research, 123(2):224–227, August 1990. ISSN 0033-7587.

Toshiaki Nakano, Ken Akamatsu, Masataka Tsuda, Ayane Tujimoto, Ryoichi Hirayama, Takeshi Hiromoto, Taro Tamada, Hiroshi Ide, and Naoya Shikazono. Formation of clustered DNA damage in vivo upon irradiation with ionizing radiation: Visualization and analysis with atomic force microscopy. Proceedings of the National Academy of Sciences, 119(13):e2119132119, March 2022. doi: 10.1073/pnas.2119132119.

G. Obe and M. Durante. DNA Double Strand Breaks and Chromosomal Aberrations. Cytogenetic and Genome Research, 128(1–3):8–16, March 2010. ISSN 1424-8581. doi: 10.1159/000303328.

Tannishtha Reya, Sean J. Morrison, Michael F. Clarke, and Irving L. Weissman. Stem cells, cancer, and cancer stem cells. Nature, 414(6859):105–111, November 2001. ISSN 1476-4687. doi: 10.1038/35102167.

Michelle Ricoul, Tamizh Selvan Gnana Sekaran, Patricia Brochard, Cecile Herate, and Laure Sabatier. γ-H2AX Foci Persistence at Chromosome Break Suggests Slow and Faithful Repair Phases Restoring Chromosome Integrity. Cancers, 11(9):1397, September 2019. ISSN 2072-6694. doi: 10.3390/cancers11091397.

Lydia Robert, Jean Ollion, Jerome Robert, Xiaohu Song, Ivan Matic, and Marina Elez. Mutation dynamics and fitness effects followed in single cells. Science, 359(6381):1283–1286, March 2018. doi: 10.1126/science.aan0797.

Evelyne Sage and Naoya Shikazono. Radiation-induced clustered DNA lesions: Repair and mutagenesis. Free Radical Biology and Medicine, 107:125–135, June 2017. ISSN 0891-5849. doi: 10.1016/j.freeradbiomed.2016.12.008.

Wook-Geun Shin, Jose Ramos-Mendez, Ngoc Hoang Tran, Shogo Okada, Yann Perrot, Carmen Villagrasa, and Sebastien Incerti. Geant4-DNA simulation of the pre-chemical stage of water radiolysis and its impact on initial radiochemical yields. Physica medica: PM: an international journal devoted to the applications of physics to medicine and biology: official journal of the Italian Association of Biomedical Physics, 88:86–90, August 2021. ISSN 1724-191X. doi: 10.1016/j.ejmp.2021.05.029.

M. Tomezak, C. Abbadie, E. Lartigau, and F. Cleri. A biophysical model of cell evolution after cytotoxic treatments: Damage, repair and cell response. Journal of Theoretical Biology, 389:146–158, January 2016. ISSN 00225193. doi: 10.1016/j.jtbi.2015.10.017.

Murat Tuğrul and Ulrich K. Steiner. Demographic consequences of damage dynamics in single-cell aging. Physical Review Research, 7(1):013327, March 2025. doi: 10.1103/PhysRevResearch.7.013327.

Michael M. Vilenchik and Alfred G. Knudson. Endogenous DNA double-strand breaks: Production, fidelity of repair, and induction of cancer. Proceedings of the National Academy of Sciences, 100(22): 12871–12876, October 2003. doi: 10.1073/pnas.2135498100.

C. K. Chris Wang. The progress of radiobiological models in modern radiotherapy with emphasis on the uncertainty issue. Mutation Research/Reviews in Mutation Research, 704(1):175–181, April 2010. ISSN 1383-5742. doi: 10.1016/j.mrrev.2010.02.001.

Hui Wang, Heng Jiang, Melissa Van De Gucht, and Mark De Ridder. Hypoxic Radioresistance: Can ROS Be the Key to Overcome It? Cancers, 11(1):112, January 2019. ISSN 2072-6694. doi: 10.3390/cancers11010112.

Daniel M. Weinreich, Tom Sgouros, Yevgeniy Raynes, Hlib Burtsev, Edison Chang, Sanyu Rajakumar, Ignacio G. Bravo, and Csenge Petak. Wanted: A population genetic theory of biological noise regulation. PLOS Genetics, 22(3):e1012066, March 2026. ISSN 1553-7390. doi: 10.1371/journal.pgen.1012066.

